# Molecular structure of the intact bacterial flagellar basal body

**DOI:** 10.1101/2020.12.05.413195

**Authors:** Steven Johnson, Emily J. Furlong, Justin C. Deme, Ashley L. Nord, Joseph Caesar, Fabienne F.V. Chevance, Richard M. Berry, Kelly T. Hughes, Susan M. Lea

## Abstract

Bacterial flagella self-assemble a strong, multi-component drive shaft that couples rotation in the inner membrane to the microns-long flagellar filament that powers bacterial swimming in viscous fluids. We here present structures of the intact *Salmonella* flagellar basal body, solved using cryo-electron microscopy to resolutions between 2.2 and 3.7 Å. The structures reveal molecular details of how 173 protein molecules of 13 different types assemble into a complex spanning two membranes and a cell wall. The helical drive shaft at one end is intricately interwoven with the inner membrane rotor component, and at the other end passes through a molecular bearing that is anchored in the outer membrane via interactions with the lipopolysaccharide. The *in situ* structure of a protein complex capping the drive shaft provides molecular insight into the assembly process of this molecular machine.

---

The bacterial flagellum is a fascinating molecular machine that is responsible for motility in many species, using a rotary motor to couple ion-flow across the inner membrane to rotation of a helical appendage on the surface of the cell^1–3^. The immense complexity of this organelle, both structural and functional, is such that it became a poster child for the irreducible complexity community; yet at its heart is a remarkable structure that reveals a great deal about assembly of complex objects from modular components.

The basal body of the bacterial flagellar motor is broadly composed of four structures; LP-ring, rod, MS-ring, and C-ring^4^. The MS-ring functions as a structural adaptor, interacting with the C-ring that coordinates the peripheral stators to generate torque^5,6^, and with the export gate responsible for secretion of the axial components of the rod and flagellar filaments^7^. Recent structures of the MS-ring revealed this adaptor function to be facilitated by structural complexity, with a single protein (FliF)^8^ assembling to form multiple sub-rings of variable symmetry^9^. The rod is a rigid helical assembly that transmits torque to the hook and filament and has historically been divided into two sections; proximal and distal. The proximal rod is composed of four different proteins, FliE, FlgB, FlgC and FlgF, while the distal rod is composed of multiple copies of just one protein, FlgG^10,11^. To date the only structural information regarding the rod, and its interactions with other components of the basal body, has come from a crystal structure of a FlgG domain docked into a 7 Å resolution cryo-electron microscopy (cryo-EM) map of an “poly-rod” formed by FlgG mutant variants^12^. The LP-ring is suggested to be the bushing of the motor, separating the rotating distal rod from the outer membrane and peptidoglycan layer. The L-ring, named because of its location near the lipopolysaccharide (LPS), is composed of multiple copies of FlgH and sits in the outer membrane (OM), whereas the P-ring is composed of multiple copies of FlgI and spans the peptidoglycan (PG) layer^13,14^.

Though recent advances in cryo-EM have enabled the determination of high-resolution structures of subcomplexes of the bacterial flagellar basal body, whole assembled basal body reconstructions (obtained mostly by cryo-electron tomography) have remained less detailed and the axial rod components have remained completely elusive. Here, we have used single-particle cryo-EM to determine structures of the intact flagellar basal body from the bacterium *Salmonella enterica* serovar Typhimurium. Our structures range in resolution from 2.2 to 3.7 Å, providing detailed structural information about the LP-ring, rod proteins, MS-ring and components of the export gate within the assembled basal body, as well as the interactions between these subcomplexes. We also observe, for the first time, an intact capping protein responsible for correct assembly of the axial components, thereby gaining insight into the mechanism of helical assembly.

We produced intact basal bodies from a mutant strain of *S.* Typhimurium that lacked the hook protein that connects the rod to the flagellar filament and was locked in a clockwise rotation mode. Complexes were extracted from the membrane and subjected to analysis by cryo-EM (Extended Data Fig. 1). All sub-structures of the basal body were visible in the particles; however, there was significant heterogeneity in the C-ring, both in terms of occupancy and conformation, so this region was excluded from further analysis. Initially, clear detail was only observed in 2D images in the region of the LP-ring, and reconstruction of this region with a variety of symmetries led to a clear solution with C26 symmetry. Classification and focused refinement produced a 2.2 Å map that allowed *de novo* building of 26 copies each of FlgH and FlgI (Fig. 1, Extended Data Fig. 2). The FlgH monomer is highly extended, centered on a small β-barrel from which extend complex loop insertions, with one such insertion extending almost 100 Å at an ~ 50° angle to the vertical axis (Fig. 1d,e). Lateral association of FlgH monomers creates a large β-barrel with an inner diameter of ~ 140 Å, the surface of which is braced by two helical bands at the position of the OM, and a shorter, but larger diameter β-barrel at the base. The N-terminus of FlgH (residues 22-69) is extended but ordered, snaking across the complex and contacting multiple subunits until terminating at the lipidated N-terminal Cys residue that associates with the OM. This N-terminal extension also serves the main contact point for the P-ring in the periplasm. The FlgI monomer is formed from 4 domains with complex, interlacing topology (Fig. 1d,e). The C-terminal domain consists of a pair of helices surrounded by paired β-strands on either side, and assembly of the P-ring brings together pairs of strands from neighboring FlgI monomers to create a four stranded sheet.

**Figure 1.**
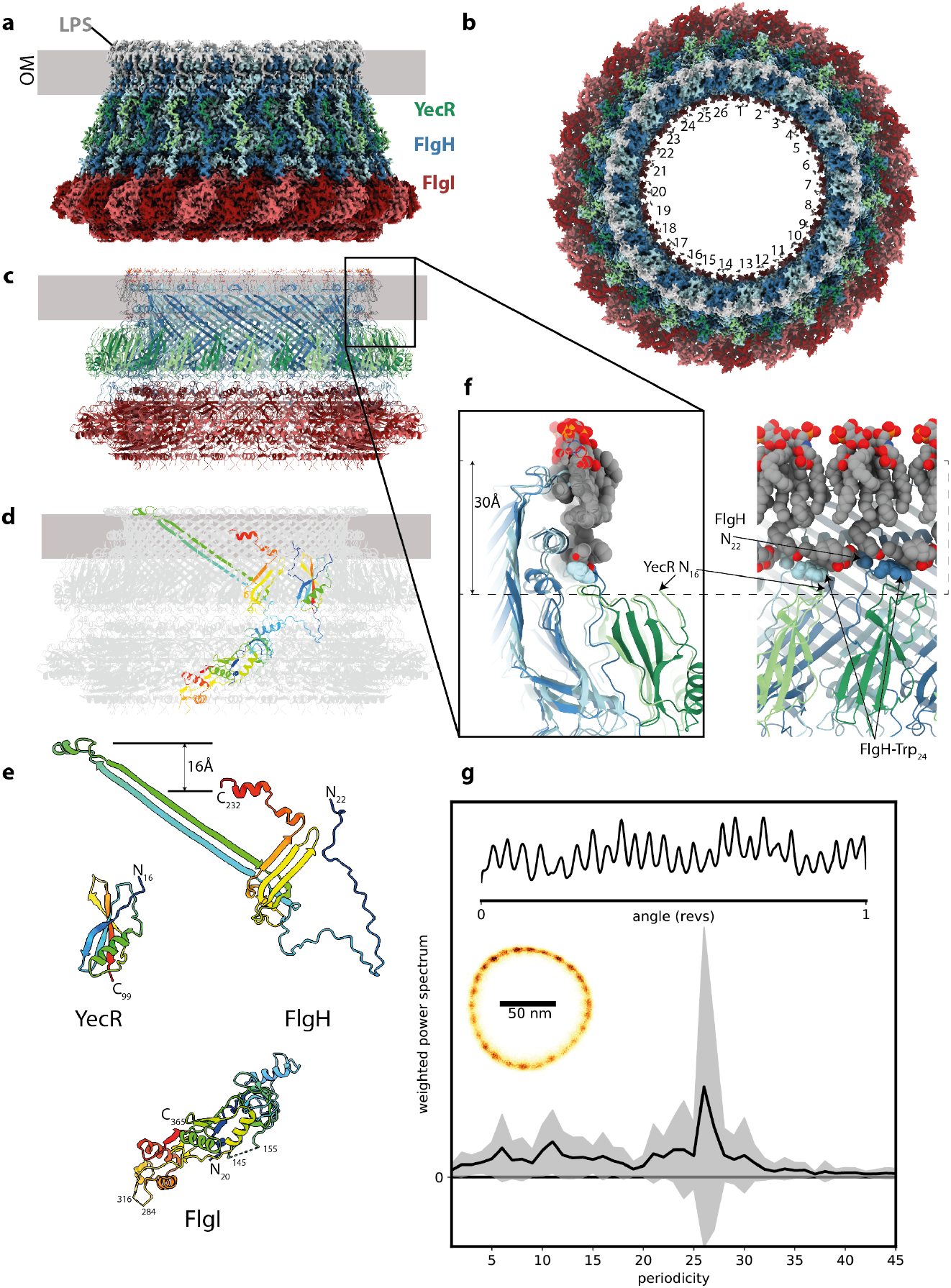
The flagellar outer membrane bushing is a lipid anchored 26mer. **a**, Side view of the 2.2 Å experimental cryo-EM volume for the LP ring bushing colored FlgI – red tones, FlgH – blue tones, YecR – green tones, lipopolysaccharide and lipidation on N-terminus of FlgH – grey. The approximate position of the outer membrane is indicated by the grey bar based on location of the detergent micelle at low contour levels. **b**, Top view of the cryo-EM volume showing the 26-fold symmetry colored as in (a). **c**, Cartoon representation of atomic model for LP ring colored as in (a) and **(d)**colored grey with the exception of a single copy of each chain which is rainbow colored from blue to red N to C terminus. **e**, View of each unique protein chain in the LP ring rainbow colored. **f**, Lipid interactions at top of LP ring assembly the two panels are related by a 90 degree rotation and show the protein in cartoon representation (colored as in (a) and the associated lipopolysaccharide and lipidated N-terminus as VDW spheres (colored C-grey, O-red, N-blue). The approximate height of a generic membrane is indicated. **g**, 26-fold periodicity in rotation of chimeric flagellar motors in *E. coli*. Left inset: 2Dhistogram of a 1 s recording of the position of a 100 nm gold nanoparticle attached to the hook of a single flagellar motor rotating at ~ 150 Hz; Top inset: corresponding kernel density distribution of the motor rotation angle. Main panel: the mean +/− S.D. (line and shading, respectively) of the weighted power spectrum of 27 distributions similar to the top inset, each from a 1-2 s recording of different cells, where the sodium motive force ranged from 54 - 187 mV and speeds ranged from ~ 2-500 Hz.

The high resolution of the LP-ring map reveals details of the interactions of the bushing complex with the OM. Associated with the top most helix of each copy of FlgH is a density that corresponds to the Lipid A portion of the LPS (Fig. 1f), with extensive hydrophobic and charged interactions anchoring it in place in the outer leaflet of the OM. Strong densities are also observed for the 3 lipid moieties of the N-terminal cysteine, with the diacylglycerol attached to the sidechain running laterally around the structure and packing against hydrophobic residues of the FlgH (Extended Data Fig. 2). Together the LPS layer and lipidation form a hydrophobic band around the L-ring that is thinner than a canonical bilayer. An additional layer of protein density was also observed outside of the L-ring, corresponding to a single domain protein bound to each FlgH. A combination of proteomic analysis and density-based sequencing identified this protein as YecR, a lipidated OM protein known to be regulated by the flagellar master regulator^15^. Notably, the position of the N-terminal cysteine in YecR is more consistent with the height of a canonical membrane bilayer (Fig. 1f), suggesting it may facilitate assembly by regulating remodeling of lipids around the unusual L-ring structure.

The LP-ring is the only flagellar component observed to have 26-fold symmetry, and therefore must be responsible for the 26 steps previously observed in partially de-energized or damaged flagella rotation^16^. These steps can also be observed in fully energized, undamaged flagellar motors (Fig. 1g). The original hypothesis that steps correspond to dynamic torque-generating interactions between stator and C-ring is inconsistent with the 34-fold symmetry of the C-ring^17–19^. Instead, these steps indicate a static 26-fold interaction potential in the bearing between the rod and LP-ring^16,20^. That this periodicity is unrelated to torque generation is further supported by observation that the phase of the 26-fold symmetry component does not change with discrete speed changes (Extended Data Fig. 3, Supplementary Fig. 1), which are signatures of structural dynamics in the stator^21^.

The high-resolution LP-ring structure was subsequently used as a reference for refinement of the full basal body without symmetry imposed. This produced a 3.7 Å C1 map with clear density for the entire axial rod down to the export gate. Focused refinement of the rod portion produced a 3.2 Å map that permitted docking and re-modeling of an export gate model and *de novo* building of each of the proximal and distal rod components (Fig. 2a,b). As previously observed in the related Type III Secretion injectisomes, the periplasmic end of the export gate flexes open relative to its isolated structure in order to nucleate the helical rod (Fig. 2c)^22^. Despite no detectable sequence identity, the FliE protein is observed to be a direct structural homologue of the SctI, with the C-terminal helix packing into the export gate template and the preceding helix folding back to pack into the crenelations of the FliP (Extended Data Fig. 4). Like the SctI sub-structure, the first copy of the FliE is only ordered in the C-terminal helix, with the N-terminus of FliR taking the space the other helix would occupy (Fig. 2d, Extended Data Fig. 4). From this point, the flagellar structure diverges from the injectisome, with sequential helical addition of 5 copies of FlgB, 6 copies of FlgC, 5 copies of FlgF and finally 24 copies of FlgG (Fig. 2d). The packing of the rod components thereby follows the non-integer helical symmetry (5.5 subunits/turn) using alternating 5:6 stoichiometries. The rod subunits are built from the same core unit, whereby a C-terminal helix packs into the centre of the growing filament, with an N-terminal helix decorating the outside (Extended Data Fig. 4). But each protein has a different domain inserted between the helices to drive correct assembly of the helix. In a further level of complexity, the FlgG is observed in 3 main conformational states that build up in sequential layers, with the differences mainly in the conformation of the loop following the N-terminal helix (termed the L-stretch) (Fig. 2d). The helical parameters of the rod therefore match, and presumably seed, those of the hook and flagellar filaments. The rod helical lattice is in turn formed by the organization of the export gate, with five copies of FliP each seeding two rod/hook/filament protofilaments and FliR seeding the final filament (Extended Data Fig. 5).

**Figure 2.**
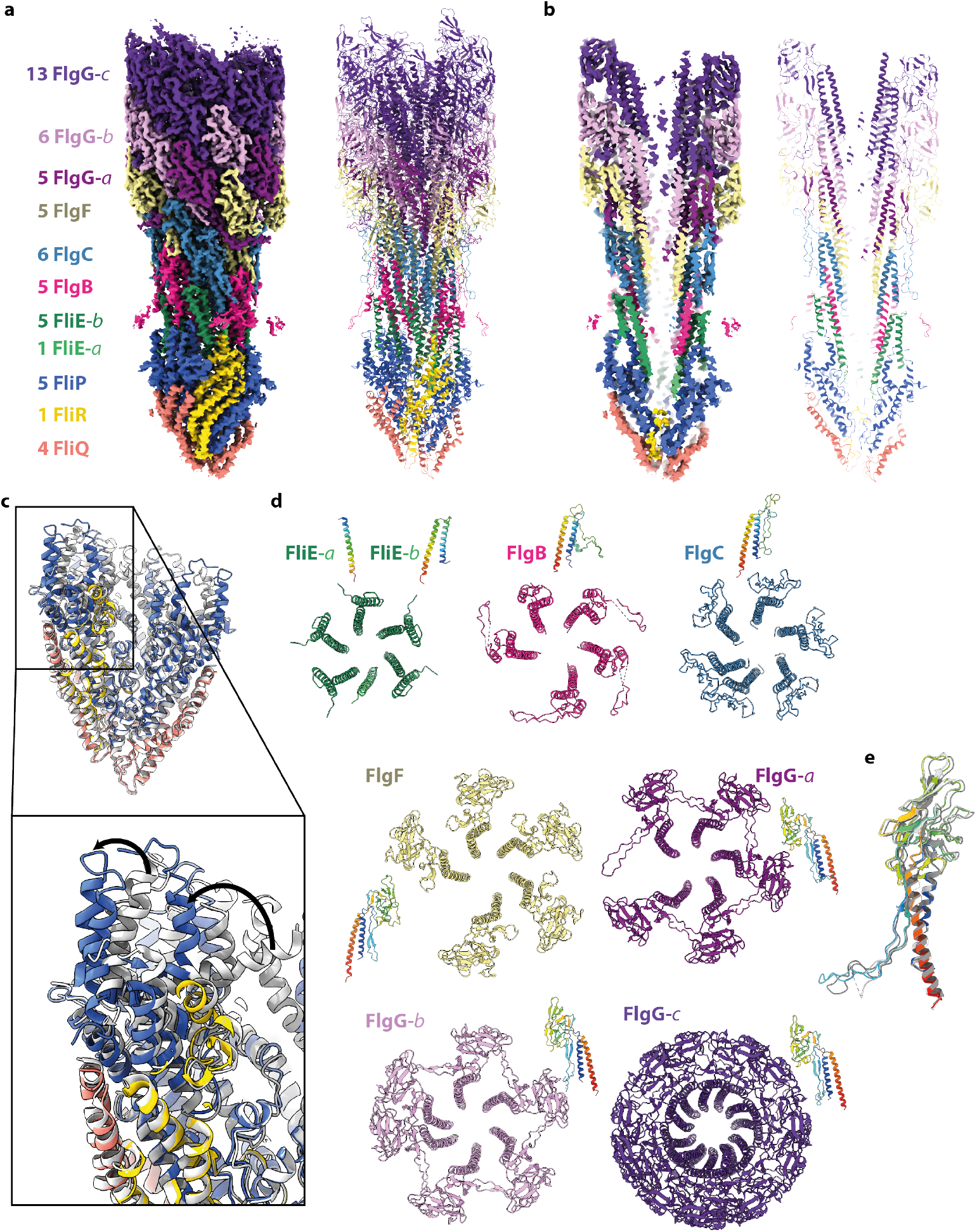
*The structure of the* in situ *flagellar axial rod*. **a**, and **b**, show the experimental cryo-EM volume (left hand side) and cartoon representation of the atomic model for the rod (right hand side) colored as defined by the key in (a). The numbers in the key indicate how many copies of each unique protein chain are observed and the -a,-b,-c postscripts indicate different conformational states of the same protein chain. The panels in (b) are a cut-through of the structure to show the central channel. **c**, The structure of the export gate as seen in the full assembly (colored as in (a)) overlaid on the structure of the isolated export gate (grey, PDB 6r69). **d**, Top views of the assemblies formed by each chain / unique conformational state of each chain are shown in cartoon representations colored as in (a) with a side view of a single subunit rainbow colored. **e**, Overlay of the three distinct conformations seen for the FlgG subunits with FlgG-a rainbow colored and FlgG-b and -c in grey.

We next analyzed contacts between the rod and the various circularly symmetric sub-structures in the basal body (Fig. 3). The LP-ring forms a bearing for the rod, with a thin, tight seal around residues 48-82 of FlgI (Fig. 3a, Extended Data Fig. 6). Opposing charges on the LP-ring seal and the rod may help to hold the bearing together (Extended Data Fig. 6), while the relatively smooth outer surface of the rod and its non-commensurate symmetry with the LP-ring allow free rotation and perhaps some degree of axial sliding. Evidence for the potential for axial sliding comes from a 3.0 Å structure of the P-ring in the absence of the L-ring (Extended Data Fig. 7), where the FlgI assembly is seen to be positioned around the lowest tier of the distal rod, closer to the MS-ring than in the final assembly. This structure also points to an important role for the membrane inserted/tethered L-ring in maintaining the P-ring bearing at the correct height relative to the outer membrane, and is consistent with earlier studies of the roles of the rod in LP-ring assembly stages^23–25^. While the mismatched symmetries of the P-ring and the rod act to minimize friction in the bearing, the C1 structure is capable of producing the 26-fold interaction potential that is observed (Fig. 1g). For example, Glu203 on sequential 1-start helix copies of FlgG forms salt bridges with Lys82 and Lys114 on two copies of FlgI, providing a local barrier to rotation on one side of the bearing. Once the torque overcomes this barrier, new salt bridges involving the same FlgG residues will form with the next set of FlgI subunits after a 13.8° rotation.

**Figure 3.**
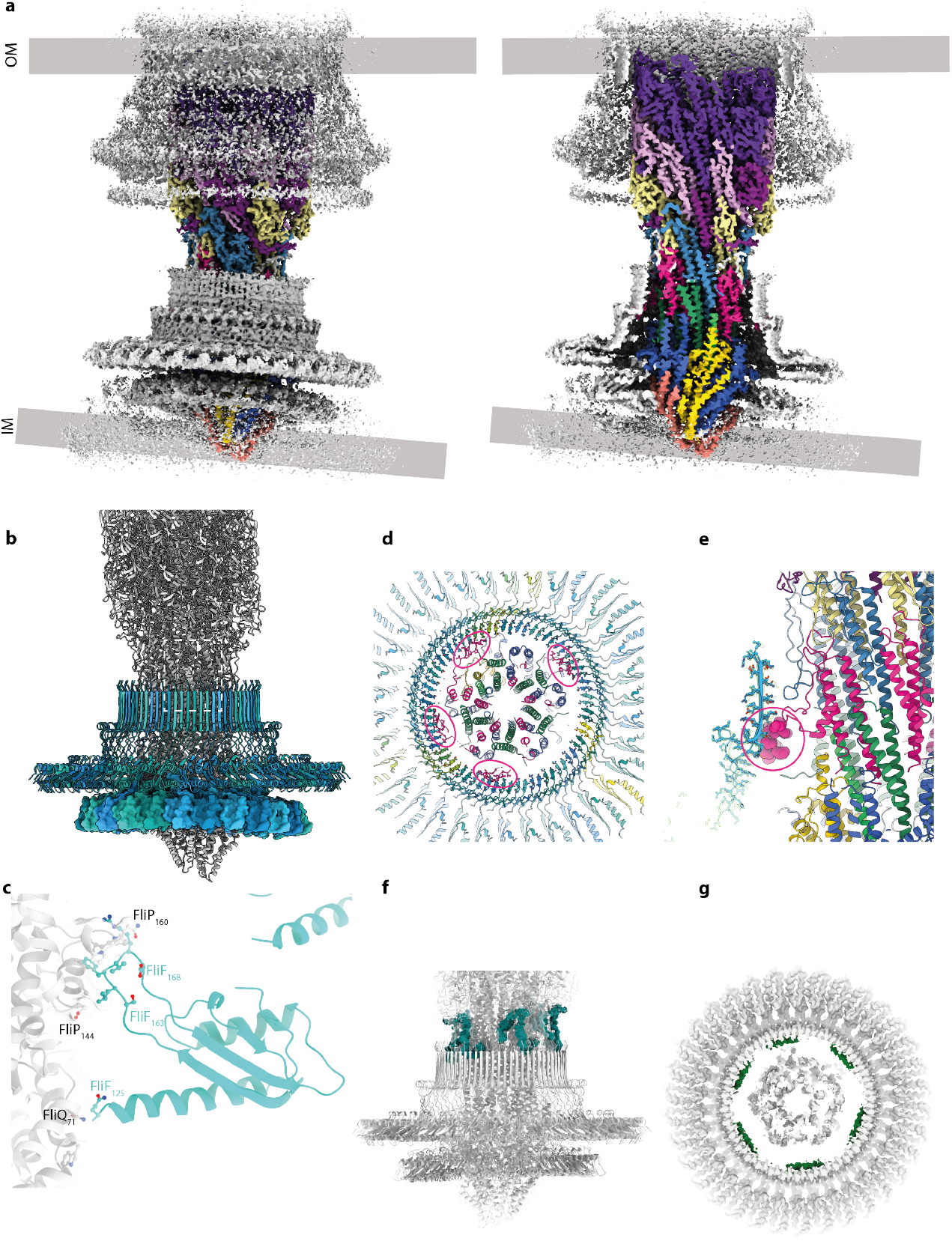
Interactions of the rod and export gate with the OM and IM ring structures. **a**, Overview of the C1 volume obtained by focusing alignments on the axial rod components. The volume is colored grey with the exception of the rod components which are colored as in Figure 2(a) on right hand panel shows a cutaway of the left hand panel. The mis-alignment of components in the inner and outer membranes is clear in this view. **b**, Cartoon of the assembly focusing on the inner membrane components with the 34mer FliF colored in blue shades. The 34-fold RBM3 ring of FliF (blue shade cartoon) is set horizontally revealing the distortion of the 23-mer RBM2 ring (surface representation) driven by contacts between the circular RBM2 oligomer and the helical export gate shown in closeup for one copy in (**c**). Other contacts between FliF and the axial components are shown by coloring the experimental volume according to the putative component contributing the contact. **d and e**, Interaction between FlgB and the inside of the FliF RBM3/ß-collar.. **f**, Putative FliF loop (teal) reaching around top of FliF assembly to contact axial rod at level of the FlgB/FlgC/FlgF interfaces. **g**, Putative FliE helix (pink) contacting base of RBM3 ß-collar.

At the proximal rod end there are myriad contacts between different chains of the rod/export gate and the MS-ring that house them, consistent with the idea that the rod must be stably attached to the rapidly rotating motor. Focused refinements around the MS-ring revealed many key details in these regions. Firstly, the FliF is observed to be a 34-mer, with the RBM3/β-collar region displaying true C34 symmetry (Fig. 3b). Contacts in this region extend in both directions, with FliF interdigitating between rod subunits, and rod subunits packing against the inside of the MS-ring β-collar (Fig. 3d,e,f,g). While the FliF densities are not sufficiently connected to unambiguously assign sequence, they clearly extend from the top of the β-collar and slot into gaps between FlgB, FlgC and FlgF, with the flexibility allowing different heights to be reached in the helix (Fig. 3f). Further down the rod, a long insertion in the FlgB (residues 58-82) packs against the midpoint of the FliF β-collar, with an interaction involving strongly co-varying residues including His72 on FlgB and Asn365 on FliF (Fig. 3d,e). Here flexibility in the linkers of the insertion allow the FlgB to pack with circular symmetry despite the helical symmetry of the FlgB core. There is also a further layer of contacts at the base of the β-collar. The six densities observed also follow the circular symmetry and look helical (Fig. 3g). These likely correspond to the predicted N-terminal helices of FliE, but again cannot be assigned in the current density. Finally, the export gate packs in the RBM2inner ring of the MS-ring. Here the RBM2inner ring is observed to contain 23 copies of RBM2, in contrast to the 22 copies observed in the isolated 34-fold MS-ring. This is analogous to equivalent region in the injectisome, where isolated structures lacking export gate contain one fewer copy of the SctJ protein^26^. There is a huge degree of flexibility in this region of the structure, with the RBM2inner ring molding itself around the helical export gate to form a highly distorted ring, unlike the rigid RBM3 ring above (Fig. 3b). Furthermore, the main loop (residues 158-173) responsible for holding the export gate also displays structural plasticity to accommodate the very different portions of structure it packs against (Fig. 3c). No evidence is seen in the maps for the FlhB component of the export gate, but this has previously been shown to be sensitive to the detergents used in extraction^27,28^. It is also possible that the detergent/lysozyme extraction process, and removal of the constraints imposed by the two membranes and peptidoglycan layer, has led to the non-coaxial relationship between the rod and the MS-ring observed.

The basal bodies characterized here are purified from cells unable to synthesize the hook protein and hence the assembly process is stalled after rod completion. The next stage in flagellar assembly would be construction of the hook itself, a process that is proposed to be catalyzed by the hook capping protein, FlgD^29^. Analysis of the end of our rod map revealed clear helical densities in the center of the completed rod that were not consistent with extra rod subunits. Further classification and focused refinement of the distal end of the rod resolved this region into five helical hairpins with clear sidechain density (local resolution of 2.9 Å) that follow the helical pitch of the rod, connected to a lower resolution (local resolution of > 7 Å) pentameric structure (Extended Data Fig. 8). *De novo* building of the helices, and docking of a crystal structure of FlgD (6iee) into the lower resolution density, allowed us to construct a model for a full length pentameric FlgD cap (Fig 4a,b,c). Residues 30-70 of FlgD are seen to form a helical hairpin (Fig. 4a), a previously unpredicted feature, though consistent with co-evolution data (Supplementary Fig. 2).

**Figure 4.**
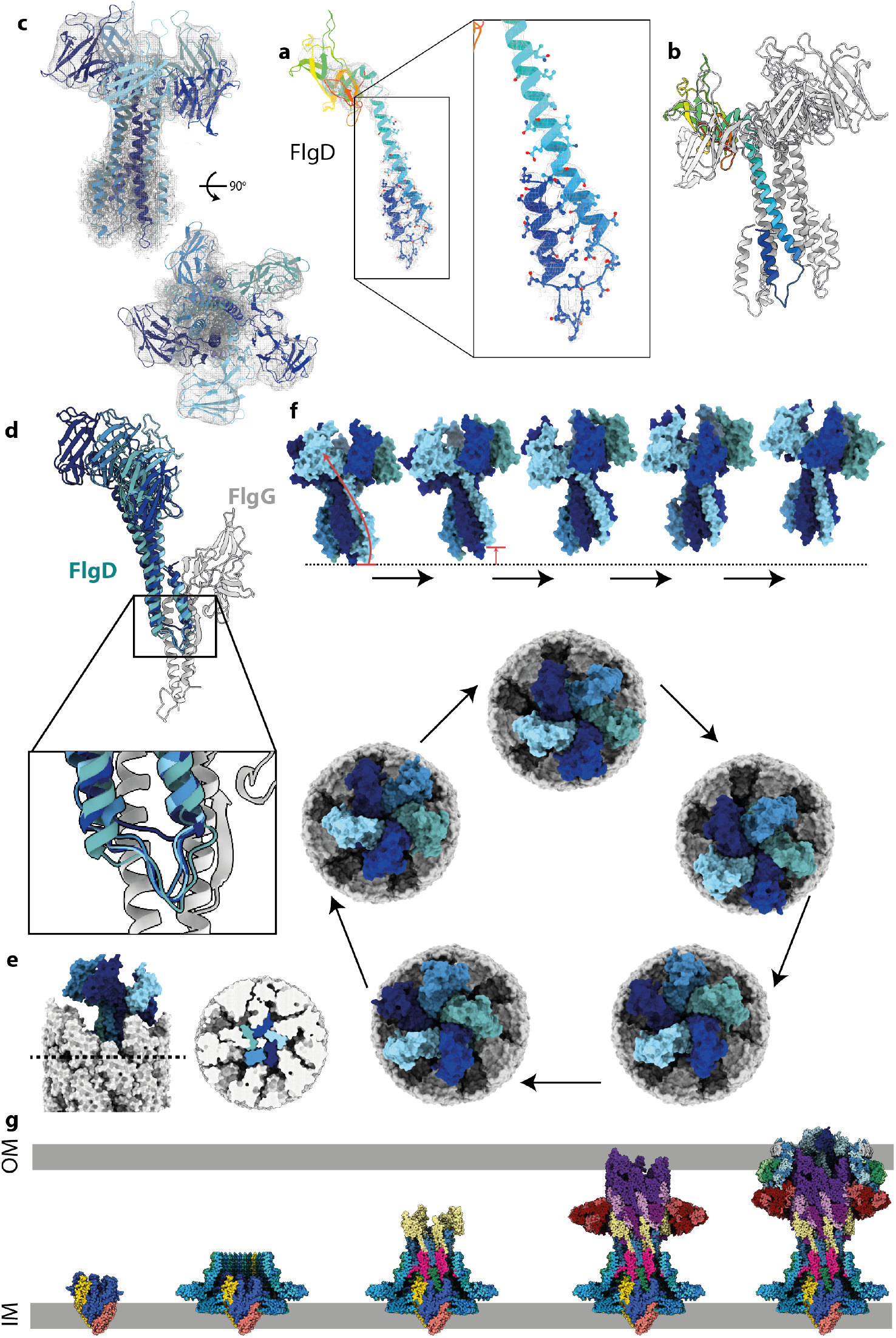
*The structure of the* in situ *hook cap complex (FlgD)* **a**, Monomer from the cap complex seen at the tip of the axial component in the Local Resolution filtered density (grey mesh) reveals the N-terminus of FlgD forms an unexpected helical pair. **b**, Pentamer of FlgD, with one copy represented as a rainbow. **c**, FlgD pentamer, colored by chain, shown in the experimental local resolution volume (grey mesh). **(d)**Each copy of FlgD primarily interacts with one copy of FlgG via stacking of the N-terminal helix pair against the FlgG N/C-terminal helices. This interaction differs for each copy of FlgD as shown in the overlay of five FlgD-FlgG pairs. One copy (navy blue) sits particularly high against its cognate FlgG, showing the FlgD helices can slip vertically with respect to FlgG. **e**, Surface representation of the tip of the axial components with the cap (cap – blue shades, FlgG – grey) *in situ* shows that the top sits at an angle to the rod axis, right hand panel shows a cut-through at the level indicated in the left-hand panel allowing visualisation of the tight packing of the FlgD blades created by the paired helices at the N-terminus. **(f)** Proposed motions involved in FlgD cap catalysed assembly mechanism. The cap is shown as a surface, colored as in part (c). Top panel shows a side view of four sequential cap movements, with the rod removed for clarity. In the first transition, the movement of the lowest copy (light blue) to become the highest copy is illustrated with an arrow. The bottom panel shows a top down view of the cap on the rod, showing a full revolution of the cap around the rod, sequentially opening up rod/hook subunit binding sites, without any rotation of the cap structure on its axis. **(g)** Assembly steps of the flagellar basal body shown using the sub-structure solved in this study. Export gate nucleates correct assembly of the MS-ring and seeds the protofilaments of the rod. Rod growth to the point of FlgG allows assembly of the P-ring around the IM-proximal end of the distal rod, which can then act as an assembly point for the H-ring. OM-tethering of the H-ring pulls the P-ring up to the appropriate height on the distal rod. Finally, FlgD assembly on the tip of the distal rod permits correct insertion of hook subunits via a stepped revolution mechanism.

The more N-terminal helix runs down into the rod structure and mimics the positioning and orientation of a rod subunit, before bending back so the more C-terminal helix of the hairpin can assemble into a 5 helix bundle with the other 4 copies (Fig. 4b). Overlay of the 5 monomers reveals a remarkable amount of flexibility in the structure (Fig. 4d), with different degrees of hinging between the 2 helices to accommodate linking the circularly symmetric C-terminal domain structure with the hairpins that follow the helical rise of the rod.

Analysis of the *in situ* cap structure provides key insights into the mechanism of cap function and filament assembly. The “resting” cap structure observed here completely blocks any exit points from the rod filament, requiring structural rearrangement to allow secretion of the next component (Fig 4e). Previous models of filament cap function have invoked rotation of the cap structures to open up the next filament subunit binding site^30,31^. However, our structure reveals that the helical hairpins are tucked deep enough into the rod interfaces to sterically prevent such a rotation (Fig. 4e). It is notable that the lowest monomer in the pentamer, which is the one facing the space for a new filament subunit, does not completely follow the helical pitch, but is sitting high as if primed for an insertion event (Fig. 4d). We therefore propose that emergence of a new filament subunit pushes this copy of FlgD in an upwards motion until it clicks into a new position at the top of the pentamer (Fig. 4f). This motion simultaneously opens up enough space to assemble the next copy of the filament, and reorients the C-terminal bundle/domain in such a way that the next FlgD is pulled up and becomes primed for the insertion event (Fig. 4f, Supplementary Movie 1). When repeated, this mechanism leads to a “stepped revolution” of the cap on the top of the growing filament (Fig. 4f).

In summary, our flagellar basal body structures provide the first sub 3 Å structures of the OM LP-ring, the entire flagellar rod and an *in situ* filament cap complex, and provide insight into the ordered assembly processes of this immensely complex molecular machine (Fig 4g).

## Acknowledgments

We thank Errin Johnson and Adam Costin of the Central Oxford Structural Molecular Imaging Centre (COSMIC) for assistance with data collection; Hans Elmlund (Monash) for access to SIMPLE code ahead of release; Nigel Moriarty, Paul Emsley and Garib Murshudov for help with modelling lipid ligands. We acknowledge use of the University of St Andrews BSRC Mass Spectrometry facility.

## Author contributions

S.J. and S.M.L. designed the project, interpreted the EM data and built atomic models. E.J.F. optimized the preparation of the basal body samples, prepared all samples and made all EM grids. J.C.D screened EM grids and together with S.M.L. collected the EM data. J.C. assisted with EM data processing. A.L.N. and R.M.B. collected and interpreted data from the motor rotation experiments. F.F.V.C. and K.T.H. created the bacterial strain used for basal body preparation. S.J., S.M.L and E.J.F. contributed to writing the first draft of the manuscript and all authors commented on manuscript drafts.

## Materials and Methods

### Materials

All chemicals were obtained from Sigma-Aldrich unless otherwise stated.

### Bacterial strains

Basal bodies were purified from *S.* Typhimurium strain TH25631 (Δ*flgE7659 flhD8070 flhC8092 fliA5225* Δ*fliB-T7771 fliG8835*(ΔPAA) Δ*rflM8403 fljB^enx^ vh2*), which expresses elevated numbers of hook-basal body structures per cell due to elevated FlhDC levels^32^. The *flhD8070*(L22H) and *flhC8092*(Q29P) were isolated as resistant to ClpXP protease degradation^33,34^. The *rflM* gene encodes a repressor of *flhDC* transcription^35^, and was deleted by recombineering using the *tetRA* cassette replacement method as described^36^. The *fliG8835* allele has a three amino acid deletion (Pro169-Ala170-Ala171) and is an extreme clockwise rotation mutant of the flagellar rotor ^37^. The *fliG8835* 3-amino acid deletion allele was constructed in TH25631 by first replacing the Pro169-Ala170-Ala171 region of *fliG* with a *tetRA* cassette followed by replacement of the *tetRA* cassette by oligonucleotide targeted replacement of the *tetRA* cassette that leaves the 3-amino acid deletion as described^36^. The strain TH26465 (Δ*yecR*∷*tetRA*) was generated by recombineering.

*E. coli* strain YS34 (ΔCheY, fliC∷Tn10, ΔpilA, ΔmotAmotB), transformed with plasmid pYS13 (pomApotB^38^, IPTG inducible, chloramphenicol resistance, pMMB206 derivative), was used for gold nanoparticle motor tracking experiments. These cells were grown from frozen aliquots in 5 ml of tryptone broth (1 % Bacto tryptone (Difco), 0.5 % NaCl) at 30°C with appropriate antibiotics and inducers (chloramphenicol, 25 μg/ml, IPTG, 30 μM) for 4.5-5.5 hours.

### Basal body purification

The purification of basal bodies from *S.* Typhimurium strain TH25631 was based on protocols published previously^39–41^. The strain was plated from glycerol stocks on LB agar, then colonies were picked and grown overnight at 37°C in LB medium. Eight liters of LB medium, in 2.5 L baffled shaker flasks, was inoculated with the overnight culture (4 mL/L) and then incubated at 37°C, 200 rpm until an OD_600_ of 0.9-1, was reached (approximately 3.5 h). Cells were harvested by centrifugation at 4,000g for 15 min, 4°C. Cell pellets were resuspended in 160 mL of ice-cold sucrose solution (0.5 M Sucrose, 0.15 M Trizma base (unaltered pH)) and while the cell resuspension was stirred at 4°C, lysozyme and EDTA pH 4.7 were slowly added (over 5 min) to final concentrations of 0.1 mg/ml and 2 mM, respectively. After 5 min of stirring at 4°C, the resuspension was moved to room temperature and stirred slowly for 1 h to allow the formation of spheroplasts. To lyse the cells, Triton X-100 was added to a final concentration of 1% v/v, and the solution was stirred rapidly for 10 min, until it became translucent. To completely degrade the DNA, 2 mg of DNase I and MgSO_4_ (5 mM final concentration) were added to the lysate. After 5 min, EDTA pH 4.7 was added to a final concentration of 5 mM. The volume of the lysate was made up to 240 mL with sucrose solution, then unlysed cells and cell debris were removed by centrifugation at 15000 g for 10 min, 4°C. Supernatant was collected and centrifuged at 142,000g for 1 h, 4°C to collect basal bodies. Pellets were resuspended in 60 mL of resuspension buffer (0.1 M KCl, 0.3 M Sucrose, 0.1% v/v Triton X-100, unbuffered) and centrifuged again at 104,000g for 1 h, 4°C. The pellet was resuspended in 2 mL of TET buffer (10 mM Tris pH 8, 5 mM EDTA pH 8, 0.1% Triton X-100) then loaded onto 20-50% w/w sucrose gradients in 10 mM Tris pH 8, 5 mM EDTA pH 8, 0.03% Triton X-100, made with a BioComp Gradient Station. Sucrose gradients were centrifuged for 14 h at 60,000g, 4°C and then fractionated. Gradient fractions were analysed by SDS-PAGE and negative stain electron microscopy and those containing basal bodies were pooled and dialysed against 10 mM Tris pH 8, 5 mM EDTA pH 8, 0.03% Triton X-100 in 5 mL 300 kDa molecular weight cut-off (MWCO) Float-A-Lyzer^®^G2 Dialysis Device (Spectrum Laboratories, Inc.) overnight. Dialysed basal bodies were then concentrated to an A_280_ of 1.5, using a 300 kDa MWCO Nanosep^®^ centrifugal concentrator (PALL).

### Cryo-EM sample preparation and imaging

Cryo-EM grids were prepared using a Vitrobot Mark IV system (FEI) at a temperature of 4°C and 100% humidity. Basal body samples were applied to graphene oxide coated^42^ Quantifoil Cu 300 mesh R 2/1 grids for 60 s before being blotted for 3 s, force −5 and then plunged into liquid ethane. Data were collected in counted super-resolution mode on a Titan Krios G3 (FEI) operating at 300 kV with a BioQuantum imaging filter (Gatan) and K3 direct detection camera (Gatan) using a physical pixel size of 0.832 Å. An initial 21,102 movies were collected at a dose rate of 15.4 e-/pix/s and exposure of 2.66 s, corresponding to a total dose of 59.2 e-/Å2, followed by collection of an additional 41,178 movies at dose rate of 14.5 e-/pix/s, exposure of 2.80 s, and total dose of 58.5 e-/Å2. All movies (62,280 total) were collected over 40 fractions.

### Cryo-EM data processing

Micrographs were initially processed in real time using the SIMPLE pipeline ^43^, using SIMPLE-unblur for motion correction, SIMPLE-CTFFIND for CTF estimation and SIMPLE-picker for particle picking. Following initial 2D classification in SIMPLE to remove poor quality particles, all subsequent processing was carried out in in RELION-3.1^44^. 43843 particles from the first dataset were re-extracted and subjected to further 2D classification. C-ring occupancy of the extracted basal bodies was observed to be variable, so selected particles (27214) were re-extracted in 432 x 432 pixel boxes centered on the LP-ring. Following another round of 2D classification, the most intact 18041 particles were refined in a variety of symmetries using an initial model generated from class averages. Only C26 symmetry produced maps with sidechain detail. The particles were subjected to two rounds of CTF refinement and one round of Bayesian polishing^45^, following which 3D classification produced a class with 10898 particles that refined to 2.2 Å resolution map using gold standard refinement. The second dataset produced 155747 particles after SIMPLE 2D, and these were re-extracted in RELION3.1 and subjected to 2D classification followed by re-extraction in LP-centered 432 x 432 pixel boxes. Combination with the first dataset and further 2D classification produced a 10237 particle subset which clearly lack the L-ring. These were refined with C26 symmetry from an LP-ring initial model low pass filtered to 15 Å resolution. A single round of 3D classification produced a set of 3194 particles that refined to a 3.0 Å resolution map using gold standard refinement.

The second dataset was also selected for intact LP-ring particles, producing 54766 particles that were combined refined and subjected to CTF refinement. These particles were then combined with the 27214 particles from the first dataset. This combined particle set was re-extracted in a 768 x 768 pixel box and subjected to 2D classification, from which 71517 particles were selected. These particles were put into masked refinement in C26 with an initial model generated from the LP-ring structure and a mask around the LP-ring. The resulting refinement was used to Bayesian polish the particle set, after which refinements were attempted with no symmetry imposed (C1). Following alignment of particles with a mask around the MS-ring and proximal rod regions, a full refinement with a mask around the entire basal body produced a 3.7 Å resolution map, using gold standard refinement, with clear helical densities through the length of the whole rod. This refinement was then used as the start point for a series of C1 refinements using local searches only and masking around different portions of the structure. Refinement with a mask around the entire rod sub-structure produced a gold standard refinement with a resolution of 3.2 Å. Refinement with a mask around the LP-ring produced a gold standard refinement with a resolution of 3.6 Å. Refinement with a mask around the RBM3/collar region of the MS-ring produced gold standard refinements with resolutions of 3.5 Å in C1 and 2.8 Å in C34. Finally, two rounds of 3D classification were carried out without image alignments, using a mask around the distal rod and cap region. This produced a 29748 particle set which refined to a 3.2 Å resolution map using gold standard refinement.

### Model building and refinement

Atomic models were built using Coot^46^, guided by co-evolution analysis. FlgH, FlgI, YecR, FliE, FlgB and FlgC models were built entirely *de novo* in their highest resolution maps. FlgF and FlgG were rebuilt from the crystal structure of the FlgG domain (6jf2), with the helical portions being built *de novo*. The N-terminal helical regions of FlgD were built *de novo* and the C-terminal domain structure docked as a rigid pentamer from a crystal structure. The RBM3/collar region of FliF was docked using a previous structure (6sd4), while the 23 copies of RBM2 region were built using the monomer structure from an earlier 22mer structure (6sd5). The previously determined FliPQR structure (6r69) was docked and manipulated to fit the density. Multiple rounds of rebuilding (in both globally sharpened and local-resolution filtered maps) and real-space refinement in Phenix^47^ using secondary structure, rotamer and Ramachandran restraints yielded the final models described in Table 1. All models were validated using Molprobity^48^. Figures were prepared using UCSF ChimeraX^49^ and Pymol (The PyMOL Molecular Graphics System, Version 2.0 Schrödinger, LLC).

### Sample preparation for mass-spectrometry analysis

A sample of purified basal bodies was run briefly into a 4-20% Mini-PROTEAN^®^ TGX Stain-Free™ gel (BIO-RAD), which was then stained with Instant*Blue*™ before the gel section containing the sample was excised and sent for analysis at the BSRC Mass Spectrometry Facility (University of St Andrews).

### Evolutionary covariance analysis

Coevolutionary contacts were determined by the Gremlin web-server^50^. Searches used the Jackhmmer algorithm for multiple sequence alignment, an E-value threshold of 10^-6^ and a minimum coverage of 75%.

### Motor rotation measurements

Cells were immobilised on poly-L-lysine coated glass coverslips in custom-made flow chambers. Anti-rabbit IgG modified gold particles of 100 nm diameter (BBInternational) were attached to the hook via anti-FlgE antibody as described previously^51^. Beads were tracked using back-scattering dark-field microscopy (see below) and a CMOS camera (Photron). The acquisition rate was 109.5 kHz. Experiments were performed in motility buffer (10 mM potassium phosphate, 0.1 mM EDTA, 1-85 mM potassium chloride, pH 5.0 - 7.0) at 23°C.

### Laser back-scattering darkfield microscopy

A mirror with a 1 mm hole in the center was mounted at a 45% angle as near the objective back focal plane (BFP) as possible (M2 in Extended Data Fig. 3a). Light from a 633 nm HeNa laser (Melles Griot LHX1, 10mW) passed through the hole and was focused onto the BFP of the objective (Nikon Plain Flour 100x oil, NA 1.3-1.4). The diameter of the field of view was ∼ 1 μm. The back-scattered light was collected by the objective, and reflected to the imaging pathway by the mirror with the hole and imaged onto the CMOS camera. A high-power LED (Thorlabs 617L2) and condenser (Nikon 1.4NA) allowed for brightfield imaging. This back-scattering setup was inspired by Sowa et al^52^, though the use of a mirror with a hole instead of a rod mirror allows for less than 4% loss of back-scattered light, compared to 8%. Recordings of gold beads stuck to a coverslip (not shown) show the total noise of the system, measured in a 55kHz bandwidth to be less than 1 nm2. For a typical BFM measurement, this corresponds to an angular resolution of better than 1**°**with a time resolution of 9.1 μs.

### Angular measurements

BFM rotation analysis was performed with custom MATLAB and Python programs. The position of the bead was determined using a Gaussian Mask fitting algorithm^53^. An ellipse was fit to the (x,y) position of the bead^54^, the data was transformed to a circle and the angular position of the motor was determined. Speed was calculated as the change in angular position between each frame multiplied by the frame rate.

A gaussian kernel of 0.02 rad was used to the create kernel density plots of the angular position shown in Fig 1g and Extended Data Fig. 3f. Histograms of the angular position (1.5-degree bin width) were used to calculate the power spectral density, which was then multiplied by the periodicity to give the weighted power spectrums shown in Fig 1g and Extended Data Fig. 3g.

